# Oleate metabolism using kinetic ^13^C dilution strategy deciphered the potential role of global transcription regulator *arcA* in *Escherichia coli*

**DOI:** 10.1101/2021.06.22.449418

**Authors:** Shikha Jindal, Poonam Jyoti, K.V. Venkatesh, Shyam Kumar Masakapalli

## Abstract

Microbial metabolism of long-chain fatty acids (LCFA; > C12) is of relevance owing to their presence in various nutrient niches. Microbes have evolved to metabolize LCFA by expressing relevant genes coordinated by various transcriptional regulators. Among the global transcriptional regulators, the metabolic control conferred by *arcA* (aerobic respiration control) under a LCFA medium is lacking. This work is targeted to unravel the metabolic features of *E.coli* MG1655 and its knockout strain Δ*arcA* under oleate (C18:1) as a sole carbon source, providing novel insights into the flexibility of the global regulators in maintaining the cellular physiology. Owing to the availability and cost of stable isotope LCFA tracers, we adopted a novel kinetic ^13^C dilution strategy. This allowed us to quantify the ^13^C dilution rates in the amino acids that retro-biosynthetically shed light on the central metabolic pathways in actively growing cells. Our data comprehensively mapped oleate oxidization in *E.coli* via the pathways of β-oxidation, TCA cycle, anaplerotic and gluconeogenesis. Interestingly, *arcA* knockout showed expeditious growth (~60%) along with an increased oleate utilization rate (~55%) relative to the wild-type. Δ*arcA* also exhibited higher ^13^C dilution rates (> 20%) in proteinogenic amino acids than the wild-type. Overall, the study established the de-repression effect conferred by Δ*arcA* in *E.coli*, which resulted in a phenotype with reprogrammed metabolism favouring higher oleate assimilation. The outcomes suggest rational metabolic engineering of regulators as a strategy to develop smart cells for enhanced biotransformation of LCFA. This study also opens an avenue for adopting a kinetic 13C dilution strategy to decipher the cellular metabolism of a plethora of substrates, including other LCFA in microbes.

## Introduction

Bacteria routinely encounter and adapt to a dynamic range of nutrient niche by evolving regulatory machinery that facilitates the metabolism of available substrates to yield energy for cellular growth and maintenance (Fong and Palsson, 2004; Görke and Stülke, 2008; Tramontano et al., 2018). In this paradigm, studying the metabolism of long-chain fatty acids (LCFA) in bacterial systems is quintessential as they play a crucial role in cell signalling, membrane synthesis and transcriptional controls (Nunn, 1986; Black and DiRusso, 1994). In addition, they are essential components in secondary metabolites such as biosurfactants and quorum sensing molecules (Schweizer and Hofmann, 2004). Furthermore, microbes in the mammalian gut systems encounter LCFAs in their surrounding niche, often from ketogenic diets (Campbell et al., 2003). Recent studies confirmed the presence of LCFAs like oleate (C18:1) and palmitate (C16) in wastewater and waste cooking oils from food processing industries (Jimenez-diaz et al., 2017; Awogbemi et al., 2019) with scope for further sustainable bioremediation.

Interestingly, *E.coli* assimilates a broad range of substrates, including LCFA, by expressing a series of fatty acid degradation (*fad*) genes along with the central metabolic pathway genes (Overath and Pauli, 1969; Klein et al., 1971; Pramanik et al., 1979; Fujita et al., 2007). Fatty acid degradation is under tight regulation of its global regulator, FadR, to satisfy the cell’s survival under genetic and environmental cues (DiRusso et al., 1993; Cronan and Subrahmanyam, 1998; Xu et al., 2001; Clark and Cronan, 2005). In the presence of LCFA, its acyl-CoA chain de-represses the catabolic pathway of *fad* genes by binding at the site of FadR regulon and gets released from the promoter region (Henry and Cronan Jr, 1991; DiRusso et al., 1992; Black and DiRusso, 1994; Cronan and Subrahmanyam, 1998; DiRusso and Knudsen, 2001).

Global transcriptional regulator, *arcA* (anoxic redox regulation) represses all sets of *fad* genes (*fadL, fadD, fadE, fadH, fadBA*) involved in transportation, activation and oxidation of FA pathway and central carbon pathway genes under glucose aerobic condition (Iuchi and Lin, 1988; Iuchi et al., 1989; Iuchi and Lin, 1991; Chao et al., 1997; Perrenoud and Sauer, 2005; Cho et al., 2006; Nizam et al., 2009; Federowicz et al., 2014). However, no comprehensive studies shed light on the metabolic control conferred by *arcA* when grown on LCFA like oleate (18:1). This work is targeted to unravel the metabolic features of *E.coli* MG1655 and knockout strain Δ*arcA* in combating non-native carbon source oleate aerobically. Stable isotopic labelling experiments with ^13^C substrates have been widely employed to map the metabolic phenotypes in *E.coli*. Of recent, high-throughput analytical platform like Gas-Chromatography Mass spectroscopy are widely adopted to measure the mass isotopomer distributions with high sensitivity (Long and Antoniewicz, 2019). Given the limitation (cost and availability) of ^13^C LCFAs, we adopted a novel approach of ^13^C dilution strategy by feeding pre-cultures with [1-^13^C] glucose followed by subjecting these cells to oleate [^12^C] and kinetically tracked the isotopic dilution patterns of metabolites. The denovo biosynthesis of biomass components using ^12^C oleate over time reduces the ^13^C levels. This strategy helped to characterize the metabolic phenotypes in WT and Δ*arcA.*

## Material and Methods

### Experimental Design

This study investigates the lucrative means to tap the fatty acid metabolism in *E.coli* MG1655 and its mutant strain Δ*arcA*. Physiological features like growth rate, oleate uptake rate and biomass yield were measured to determine the phenotypic characteristics of strains. A novel approach of ^13^C dilution was employed to understand the metabolic distribution in fatty acid metabolism. The dilution patterns of valid amino acid fragments were analyzed kinetically in order to get insights into the oleate metabolism in both strains. Datasets of mass isotopomer distribution were derived from nine valid amino acids fragments (alanine, aspartate, glutamate, glycine, isoleucine, phenylalanine, serine, valine and threonine) in triplicates.

### Chemicals

Glucose, sodium oleate and minimal media components used in this study were purchased from Sigma-Aldrich and glucose tracers from Cambridge Isotope Laboratory (CIL): [1-^13^C] Glucose (99.2% ^13^C).

### Bacterial strains and growth conditions

In this study, *E.coli* K-12 MG1655 (CGSC#6300) was used as the wild-type (WT) strain along with its mutant strain, Δ*arcA*, which was constructed using a one-step inactivation protocol as shown in Supporting Material 1 (Datsenko and Wanner, 2000). This knockout strain was validated by PCR. WT and Δ*arcA* were cultivated aerobically in sodium oleate (1 g per litre) as a sole carbon source in M9 minimal media (Composition per litre of distilled water: 6 g Na_2_HPO_4_*12H_2_O, 3 g KH_2_PO_4_, 1 g NH_4_Cl, 0.5 g NaCl along with 1 g MgSO_4_ and 0.5 g CaCl_2_). 5 ml of working volume was used in the batch culture at 37°C on a rotary shaker (Eppendorf). The pH of the growth medium was set to 7 ± 0.05.

### ^13^C dilution experiment

A dilution experiment was performed to understand the pathway dynamics of oleate. First, cells were pre-cultured in 30% [1-^13^C] glucose substrate till the exponential phase. Thereafter, 0.05 OD cells were washed with buffer and transferred to the media containing unlabelled sodium oleate as a sole carbon source (Fig. 1A). Cells were harvested at different time points (0h, 4h, 6h, 8h). Further, to quantify the isotopic labelling, cells were acid hydrolyzed, derivatized and then subjected to GC-MS to obtain the labelling pattern in the proteogenic amino acids (Fig. 1B). All the labelling experiments were performed in triplicates.

**Figure 1.**
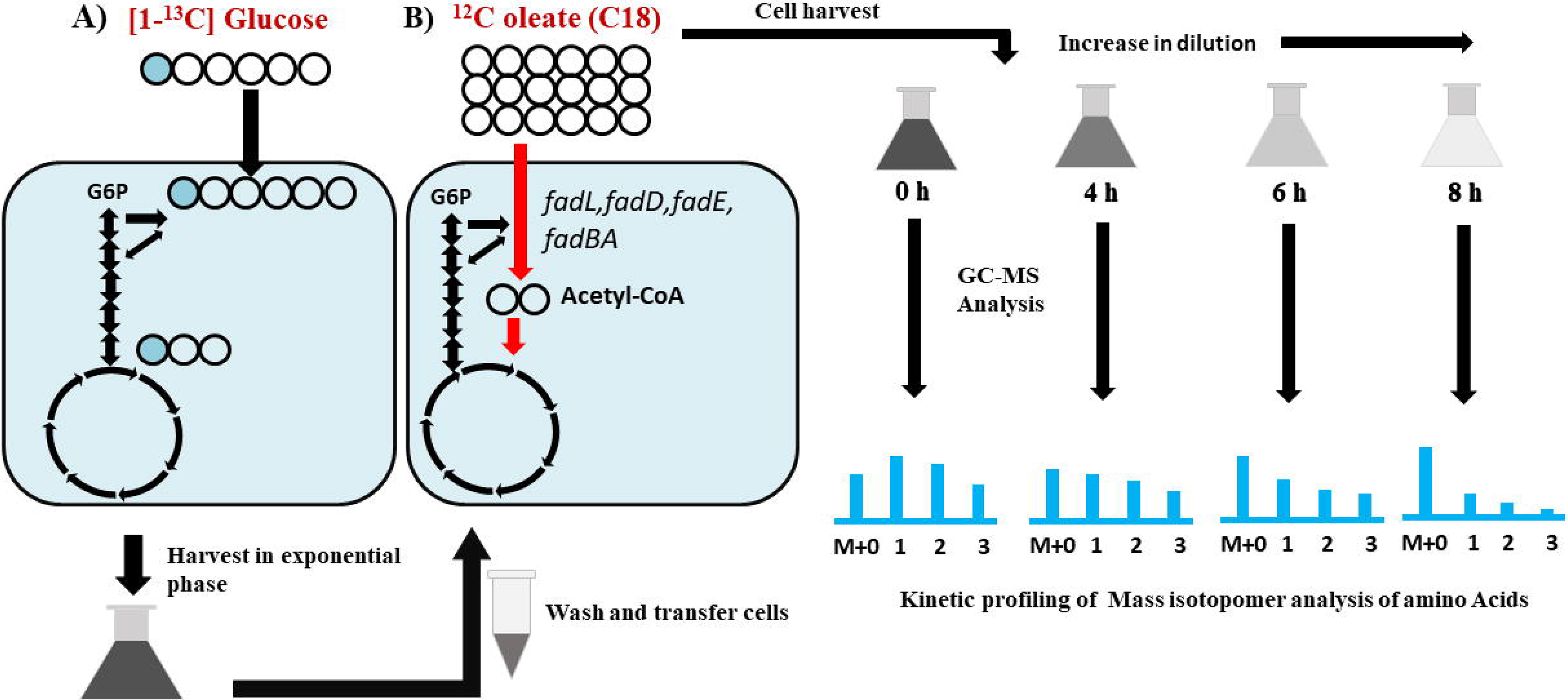
Schematic illustration of the oleate metabolism workflow in *E.coli* A) 30% [1-^13^C] Glucose was fed into the pre-culture in WT and Δ*arcA*. A) Generation of ^13^C labelled *E.coli* WT and Δ*arcA* cells using 30% [1-^13^C] Glucose B) Feeding of unlabeled oleate to the ^13^C cells and kinetic monitoring of the dilution in targeted molecules.

### Microbial growth, biomass and CHNS Elemental analysis

Cellular growth was monitored spectrophotometrically by measuring the optical density at 600 nm (OD_600_). For each strain, the growth rate was estimated based on linear regression of the natural logarithm of OD_600_ in the exponential phase versus time. Furthermore, the growth rate was utilized to calculate the oleate uptake rate and biomass yield (Chen et al., 2011). Cells were harvested and lyophilized in the exponential phase for the dry cell weight quantification. To determine the relationship between OD_600_ and dry cell weight (DCW), OD_600_ was measured at regular intervals in the log phase with its corresponding dry cell weight per litre. Therefore, a conversion coefficient of 0.42 gDCW/L/OD_600_ was calculated. For calculating the elemental analysis of carbon and nitrogen, 5 mg of lyophilized cell pellet was subjected to CHNS (Carbon Hydrogen Nitrogen Sulphur) analyzer (ThermoFinnigan).

### ^1^H-NMR analysis of Oleate

Oleate concentration was determined at an equal time interval in the media using ^1^H proton NMR (JNM-ECX500 spectrometer). The cell culture was centrifuged to remove the cell debris, and the supernatant was collected by filter sterilization. 300 μl of filter-sterilized supernatant was mixed with the 200 μl of double distilled water to dilute. Further, 100 μl of 0.1% DSS (4,4-dimethyl-4-silapentane-1-sulfonic acid) was added to each sample as an internal standard reference for determining the oleate concentration at a particular time point. The solution was gently mixed and transferred to 5 mm glass tubes for oleate quantification using JOEI-DELTA.

Each Spectra of ^1^H-NMR was recorded with 64 scans and a pulse width of 11.6 μs. The spectral intensities were calculated using the Delta software (version 5.7). The concentration of oleate was calculated at equal intervals in the exponential phase to calculate its consumption rate.

### Gas Chromatography-Mass Spectroscopy (GC-MS)

GC-MS measurements were executed on an Agilent 7890A GC system equipped with a DB-5ms capillary column (30 m, 0.25 mm, 0.25 μm-phase thickness; Agilent J&W Scientific), connected to a Waters Quattro Micro Tandem Mass Spectrometer (GC-MS/MS) operating under ionization by electron impact (EI) at 70 eV. An injection volume of 1 μl was used for all the samples to be analyzed with helium as a carrier gas at 0.6 ml/min. The initial temperature of the oven was at 120°C held for 5 min. It was followed by 4°C/min gradient upto 270°C, held for 3 min, followed by 20°C/min ramped to 320°C. The GC-MS spectra were recorded for a total run time of 50 min with a scanning range of 40-600 m/z.

### Sample Preparation, derivatization and GC-MS

Bacterial cells (2 mg dry weight) were subjected to acid hydrolysis by adding 500 μl of 6 M HCl and incubating at 95°C for 16 h (Shree and Masakapalli, 2018). 20 μl of acid hydrolysate was subjected to speed-vacuum (Thermo-315 scientific, Waltham, MA, USA) for 3 h at 35°C to obtain the dried extract. Subsequently, the dried extracts were subjected to TBDMS derivatization to detect the amino acids. Further, 30 ul of pyridine was added to each dried hydrolysate extract sample and incubated the tubes at 37°C for 30 min at 900 rpm on a thermomixer. Thereafter, 50 μl of MtBSTFA ((N-Methyl-N-(t-butyldimethylsilyl trifluoroacetamide)) + 1% t-BDMCS (N-methyl-N-(t-butyldimethylsilyl) trifluoroacetimide) + 1% t-320 (butyl-dimethylchlorosilane) was added to each sample and incubated at 60°C for 30 min at 900 rpm (thermomixer). Thereafter, samples were centrifuged at 13,000*g for 12 min, and 50 μl of supernatant was transferred to GC-MS vials for sample injection.

### GC-MS data analysis

The raw GC-MS data files were generated and the baseline correction was performed using MetAlign Software (Lommen and Kools, 2012) to evaluate the precise mass isotopomer distribution (MID) of all amino acids. We employed Agilent ChemStation software to obtain the intensity of the mass ions corresponding to each hydrolyzed amino acid fragment. Each derivatized amino acid was confirmed using the National Institute of Standards and Technology (NIST, Maryland) library. Here, data files were mass corrected of the natural abundance of ^13^C isotope present in the amino acid fragments (Masakapalli et al., 2014) and ^13^C incorporation was analyzed using an IsoCorr software (Millard et al., 2012). The corrected MIDs (Supporting Material 2) of each valid amino acid fragment (Antoniewicz et al., 2007) were subjected to enumerate the average ^13^C abundance with precision (Shree and Masakapalli, 2018).

### Statistical analysis

In each experiment, all data values are mentioned as mean ± Standard deviation (SD). The student’s t-test was performed to determine the statistical significance across WT and Δ*arcA* data in triplicates.

## Results

### Δ*arcA showed enhanced growth and oleate uptake rate*

The growth of *E.coli* MG1655 and its knockout strain Δ*arcA* were measured under oleate as the sole carbon substrate. Here, the growth rate, oleate uptake rate and biomass yield for both the strains are summarized in Fig. 2. Δ*arcA* exhibited a significantly higher growth rate (0.25 h-1) compared to WT (0.16 h-1) under oleate aerobic condition (Fig. 2A). The oleate uptake rate also exhibited a similar growth rate trend with a 50% higher uptake in Δ*arcA* than WT (Fig. 2B). Interestingly, the biomass yield of Δ*arcA* (0.82 gDCW/gOleate) was slightly better than WT (0.78 gDCW/ gOleate) (Fig. 2C). It is further noted that Δ*arcA* exhibited shortened lag phase than WT (Fig. 2D). Overall, Δ*arcA* displayed faster growth, slightly increased biomass yield inspite of higher oleate uptake rates.

**Figure 2.**
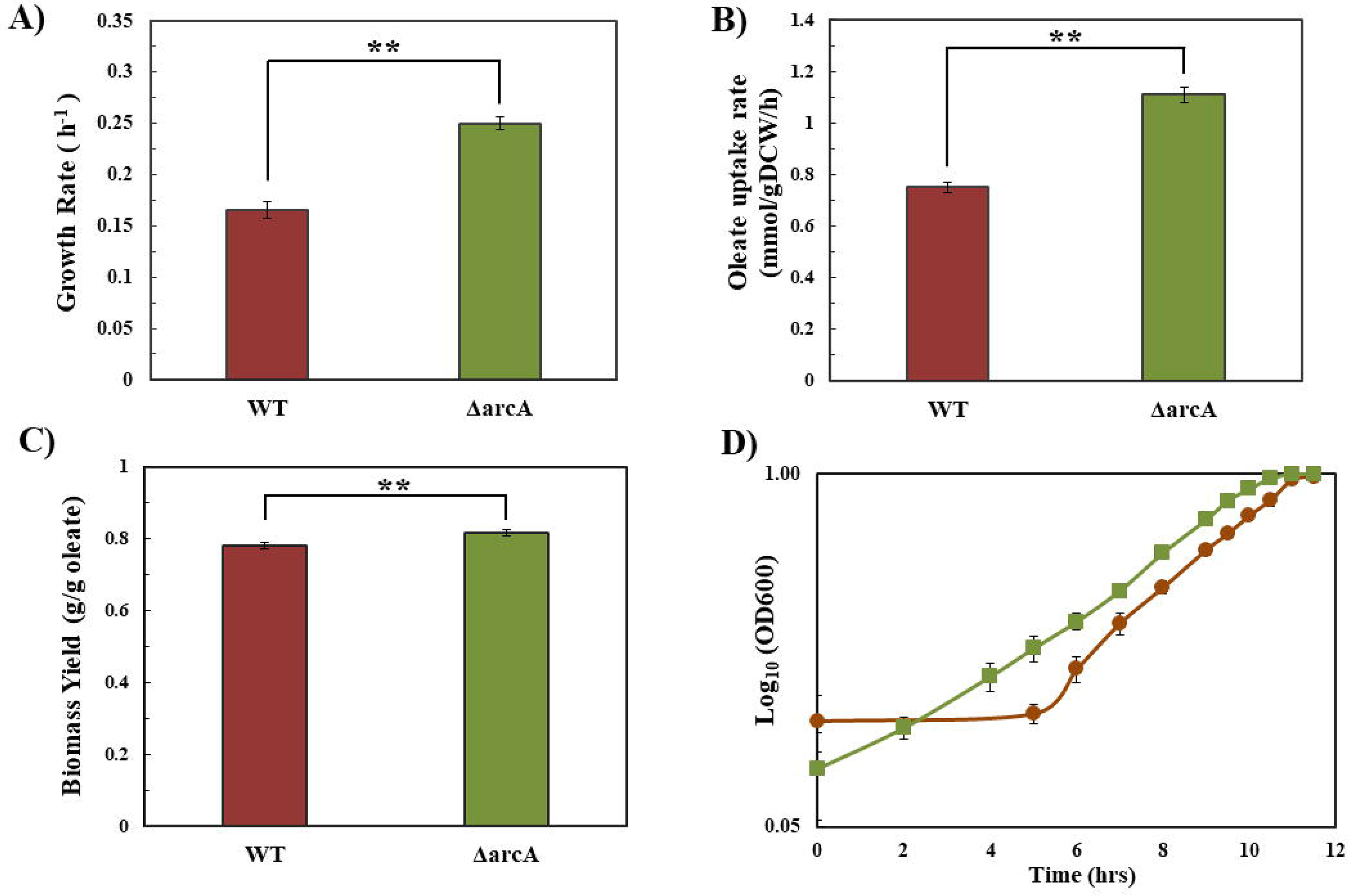
Physiological characteristics of *E.coli* wild-type and Δ*arcA* grew on oleate M9 medium. A) Growth rate B) Oleate uptake rate in mmol/gram Dry cell weight/hour measured using ^1^H NMR C) Biomass yield D) Growth profile of WT (maroon color) and Δ*arcA* (green colour). All the error bars reflect the SDs (n = 3). Two asterisks (**) denotes p < 0.01.

### Oleate is assimilated in E.coli WT via the TCA cycle, anaplerotic pathway and gluconeogenesis

We adopted a novel ^13^C dilution experiment to determine the pathway dynamics in *E.coli* MG1655 and Δ*arcA* under oleate metabolism. The ^13^C abundances in isotopomers of aminoacids decreased kinetically (from 0 h to 8 h) due to the dilution effect of oleate (Fig. 3). The unlabelled acetyl-CoA (from oleate feed) enters into the TCA cycle and contributes to the denovo biosynthesis of amino acids, thereby diluting the ^13^C levels in starting cells (Table 1; Figure 3). Fig. 3 describes the isotopomer distribution of all the valid amino acid fragments in WT where fractional enrichment in M + 0 enhanced in all sets of amino acids but declined in M+1, M+2, M+x isotopomer with time. This demonstrates the continuous influx of unlabelled oleate into the system, which furthermore elevated the unlabelled amino acid pool kinetically. One of the vital amino acids synthesized from the TCA cycle is glutamate, wherein 56% of the average ^13^C label got diluted in the early 4 h of growth in WT (Table 1, Fig. 3). By 8 h, a dilution of 91% was observed in glutamate (Table 1, Fig 3).

**Table 1.**
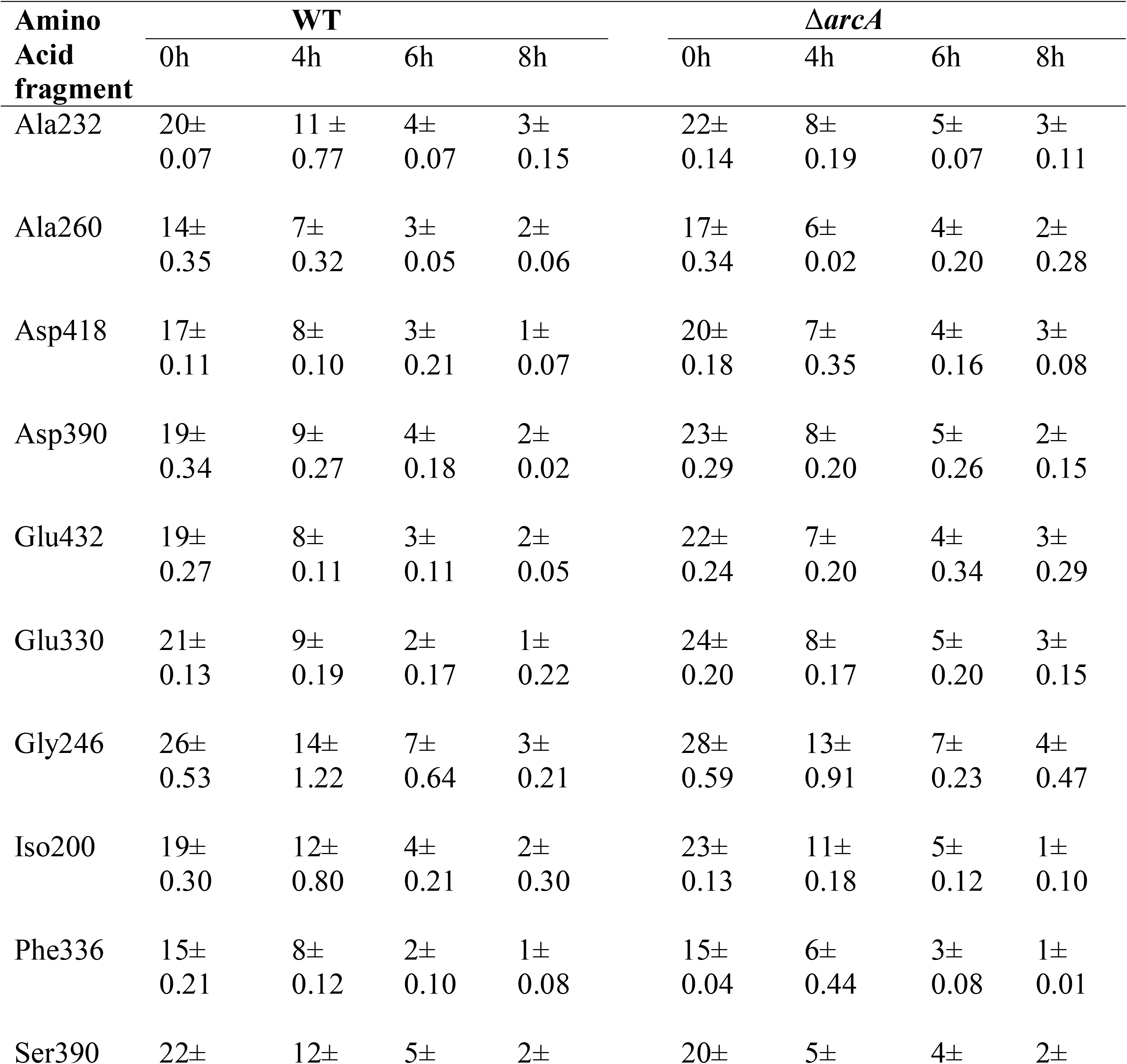
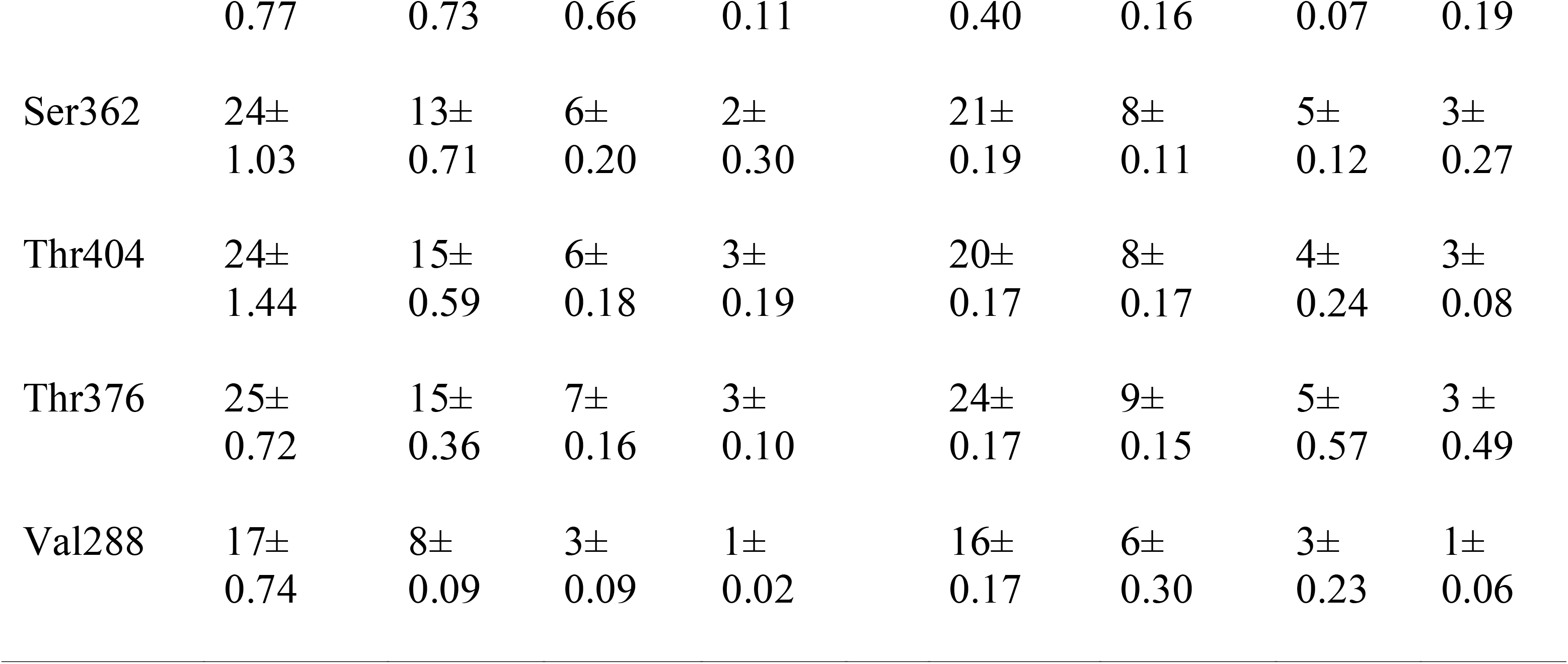
Kinetics of average ^13^C MID values (in %) decreased with time in wild-type and in Δ*arcA*. Error indicates the SDs of the mean values.

**Figure 3.**
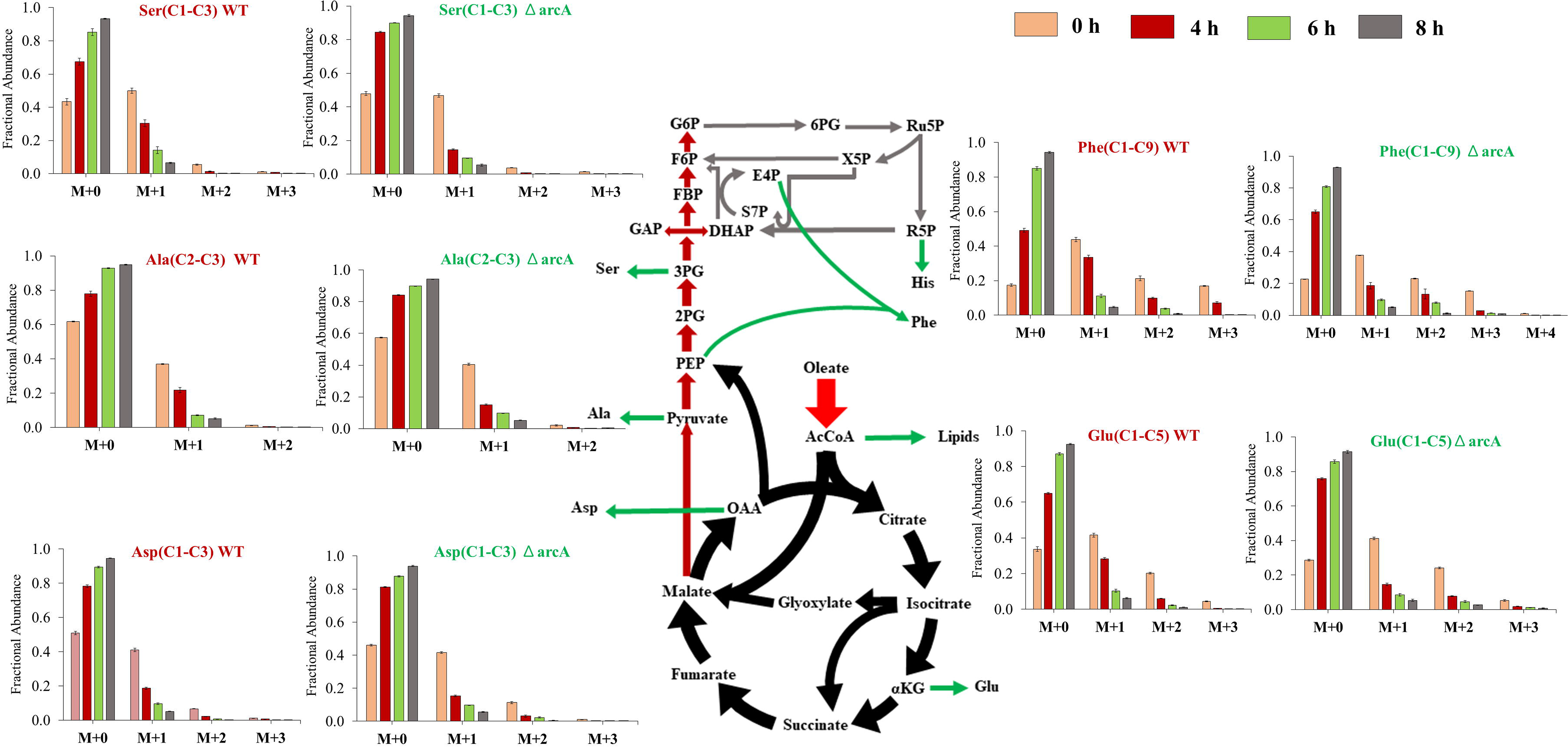
Oleate metabolism in *E.coli* and Δ*arcA* are represented by Mass isotopomer distribution in amino acids. Time course fractional abundances of representative amino acids synthesized from the TCA cycle (glutamate and aspartate), anaplerosis and gluconeogenesis (Alanine and serine) and PPP (Phenylalanine) are presented. In each amino acid fragment analyzed, (M+0, M+1…) corresponds to the MIDs. We observed that the fractional abundances varied with the incorporation of oleate into the system and from these measurements, the average ^13^C levels were derived (Table 2).

Further, oxaloacetate (OAA) synthesizes aspartate by reversible transamination reaction (Reitzer, 2004). At the 0th hour, the average ^13^C fractional enrichment in Aspartate (C2-C4) was 19% which was reduced by 56% in 4 h of growth, showing a similar trend as seen in glutamate residues (Table 2). In threonine fragments, the initial reduction in the average ^13^C label was by 38%. Thus, a high ^13^C fractional dilution rate in aspartate and glutamate fragments predicts the higher influx of oleate through the TCA cycle.

**Table 2.** The rate of average ^13^C dilution in Amino acids between growth intervals of 0 to 4 hours, 0 to 6 hours and 0 to 8 hours.

| Strains | | WT | | | $\Delta arcA$ | | |
| --- | --- | --- | --- | --- | --- | --- | --- |
| Amino acids | Carbon atoms in amino-acid | 0-4 h | 0-6h | 0-8h | 0-4h | 0-6h | 0-8h |
| Ala232 | C2-C3 | 44% | 81% | 87% | 63% | 77% | 86% |
| Ala260 | C1 to C3 | 52% | 81% | 87% | 67% | 78% | 86% |
| Asp418 | C1 to C4 | 54% | 80% | 95% | 66% | 79% | 86% |
| Asp390 | C2 to C4 | 56% | 80% | 90% | 67% | 78% | 90% |
| Glu330 | C2 to C5 | 55% | 88% | 95% | 66% | 79% | 88% |
| Gly246 | C1-C2 | 46% | 74% | 88% | 54% | 76% | 85% |
| Iso200 | C2 to C6 | 38% | 79% | 89% | 55% | 80% | 94% |
| Phe336 | C1 to C9 | 46% | 86% | 95% | 59% | 78% | 92% |
| Ser390 | C1-C3 | 45% | 76% | 89% | 72% | 82% | 90% |
| Ser362 | C2-C3 | 44% | 76% | 91% | 63% | 78% | 87% |
| Thr404 | C1 to C4 | 38% | 76% | 89% | 61% | 80% | 87% |
| Thr376 | C2 to C4 | 38% | 72% | 89% | 61% | 77% | 89% |
| Val288 | C1 to C5 | 55% | 80% | 92% | 64% | 80% | 93% |

*$\Delta arcA$ exhibited enhanced anaplerotic pathway rates qualitatively*

Malate pool derived from glyoxylate and fumarate bifurcates its fluxes to OAA and pyruvate, respectively. Pyruvate, along with aspartate, synthesizes alanine, valine and isoleucine. The ^13^C fractional dilution in the Ala(C2-C3) was 44% in 4 h and reached 87% by 8 hours. Similarly, in the Val(C1-C5), the rate of average ^13^C dilution remained similar to alanine. Interestingly, the dilution rate further decreased in the isoleucine fragment, which signifies that the rate of synthesis of alanine and valine is higher than that of isoleucine. Thus, the rate of ^13^C dilution in anaplerotic intermediates was less than the TCA cycle intermediates. Therefore, it can be predicted that the fluxes are lower in the anaplerotic pathway than in the TCA cycle.

Gluconeogenesis pathway activity would be needed to achieve biomass and energy demands. The carbon from the TCA cycle intermediates (OAA) via anaplerotic pathways feed into the gluconeogenesis leading to the production of central precursors PEP and Pyruvate. These further act as precursors for gluconeogenesis and PPP, eventually synthesizing the aromatic amino acids such as phenylalanine and tyrosine. Our results illustrated that in the initial 4 h of growth, ^13^C fractional dilution in Phe (C1-C9) reduced by 46% and further by 86% and 90% in 6 h and 8 h, respectively. Here, the ^13^C dilution rate has decreased in comparison to the anaplerotic intermediates and TCA cycle.

Further, serine and glycine are synthesized from the 3PG that represents the upper metabolic pathway of gluconeogenesis. In serine (C2-C3), the ^13^C label reduced by 44% in the initial 4 h and reduced by 89% in 8 h. Glycine (C1-C2) also exhibited a similar trend by showcasing the ^13^C dilution by 46% and 88% in 4 h and 8 h, respectively (Table 2). Overall, in *E.coli*, it is observed that the TCA cycle manifests the highest rate of dilution, followed by anaplerotic pathway and gluconeogenesis.

### *Global transcription regulator knockout* Δ*arcA exhibited enhanced TCA cycle activity*

We observed distinct oleate metabolic phenotypes in Δ*arcA* compared to the wild-type. In 4 hours, Δ*arcA* showed a higher average ^13^C isotopic dilution than the WT (Table 1). In Δ*arcA*, isotopomers (M+1, M+2) abundances reduced faster than the WT (Fig. 3). It should be noted that in Asp(C2-C4), a 67% dilution rate was observed in the initial 4 h of growth in Δ*arcA*, compared to WT (56%). The rate of average ^13^C dilution within 4 hours can capture the metabolic variations across various pathways in *E.coli* strains. In Δ*arcA*, the remarkable reduction in average ^13^C labelling was observed from 0 to 4 h, compared to WT.

The enhanced rate of ^13^C dilution was observed in glutamate, aspartate and threonine of Δ*arcA* compared to WT. In glutamate and aspartate, 1.22 folds higher ^13^C dilution rate was seen in Δ*arcA* (0 to 4 h) than the WT. In 6 h, the rate of dilution further decreased by 80% in both the strains. By the end of 8 h, only 10% of the ^13^C fractional label was retained in the pool. In threonine residues, 1.6 folds higher dilution rate was quantified in Δ*arcA* (0 to 4h) than the WT. This highlights Δ*arcA* confers de-repression in the genes controlling the oleate metabolism via the TCA cycle.

### Δ*arcA exhibited enhanced anaplerotic pathway rates qualitatively*

The MIDs of alanine can retrobiosynthetically link pyruvate, which is formed via anaplerotic pathways (OAA to Pyr, Mal to Pyr) or gluconeogenesis. In 4 hours, the ^13^C dilution rate of alanine in Δ*arcA* was 1.42 folds higher than the WT (Table 2). Thereafter, within 8 hours, the cells reached similar average ^13^C levels of ~87% in both strains. Also, the MIDs of valine can report on the Pyruvate and hence anaplerotic reaction activities. In valine fragment, 1.2 folds higher dilution rate was observed in the initial 4 h of growth in Δ*arcA* than in the WT. In 6 and 8 h, the dilution rate remained unaltered in both the strains. In Ile(C2-C6), 1.25 folds higher dilution rate was observed in Δ*arcA* from 0 to 4 h, which was quite similar in valine. The reverse trend was observed from 4 to 6 h, with an increased dilution rate in WT by 1.2 folds compared to the Δ*arcA*.

Overall, the analysis showed that alanine, valine and isoleucine exhibited a faster dilution rate in Δ*arcA* than WT. Although, the study indicates that the fluxes in the anaplerotic pathway elevated in Δ*arcA* compared to the WT. The metabolic phenotype of Δ*arcA* exhibiting higher anaplerosis as a consequence of pathway de-repression seems an important rewiring mechanism to adopt.

### Δ*arcA showed an increased rate of dilution towards gluconeogenesis*

The evident amino acid recovered from gluconeogenesis was phenylalanine, synthesized from PEP and E4P. In Phe(C1-C9), ^13^C label got diluted by 1.25 folds faster in Δ*arcA* than WT in 4 h. Between 4 to 6 h, the ^13^C label further decreased in both strains but with a higher magnitude in WT by 1.5 folds compared to Δ*arcA*. In 8 hours, the final dilution rate was > 92% in both WT and Δ*arcA*. This data corroborates that the amino acids synthesized from PEP were diluted from the convergence of two metabolites, i.e. pyruvate and oxaloacetate. In 4 h, Gly(C1-C2)246 was diluted by 48% in WT and 53% in Δ*arcA*. By the end of 8th h, the final ^13^C dilution was 88% in both strains. A decline in the label dilution rate in glycine than other pathway amino acids signifies the lower dilution rate towards the upper pathway metabolism, i.e. gluconeogenesis. In Ser(C2-C3), labelling is diluted by 43% and 63% in WT and Δ*arcA* strain, respectively, in 4 h. Thereafter, in 6 h, the dilution rate of Δ*arcA* increased by 1.37 folds compared to the WT. From G6P, the fluxes channelled towards the PP pathway for ribose production. The dilution in serine residue is indicative of the PP pathway activity in both the strains with higher activity in Δ*arcA* compared to WT. Thus, Δ*arcA* shows >25% higher dilution through gluconeogenesis and PP Pathway than WT.

## Discussion

Microbes have evolved to oxidize LCFA by expressing relevant genes coordinated by various transcriptional regulators. Among the global transcriptional regulators, the metabolic control conferred by *arcA* (aerobic respiration control) under an LCFA medium is lacking. This work is targeted to unravel the metabolic features of *E.coli* MG1655 and its knockout strain Δ*arcA* under oleate (C18:1) as a sole carbon source, providing novel insights into the flexibility of the global regulators in maintaining the cellular physiology. Growth physiology demonstrated Δ*arcA*, a faster-growing strain with its high substrate utilization rate under oleate as a carbon source.

Owing to the limitations of the availability and cost of stable isotope LCFA tracers, we adopted a novel ^13^C dilution strategy to map the pathways kinetically. This allowed us to quantify the ^13^C dilution rates in the amino acids that retro-biosynthetically shed light on the central metabolic pathways in actively growing cells. The Mass isotopic distribution in proteinogenic amino acids kinetically (0 to 8 h) accounted for the ^13^C levels and their dilution rates in Δ*arcA* and WT. In addition, the kinetics of labelled isotopomer of all amino acids was compared. Consequently, it quickly incorporated oleate in the TCA cycle, followed by an anaplerotic pathway and then towards gluconeogenesis and the PP pathway.

Interestingly, Δ*arcA* showed a greater magnitude of difference (>20%) in dilution rate of MIDs throughout the central carbon pathway, i.e. TCA cycle (Glutamate, aspartate), anaplerotic reaction (Alanine, valine) and gluconeogenesis (serine, glycine) as compared to WT. It can be inferred that in Δ*arcA*, the increased ^13^C isotopic dilution rates correlate with the de-repression effect in the oleate metabolic pathway relative to WT. Literature has reported that *arcA* is a tight regulator of the β-oxidation pathway, TCA cycle and glyoxylate shunt under glucose aerobic condition (Maloy et al., 1980; Perrenoud and Sauer, 2005; Waegeman et al., 2011). A similar effect was observed under oleate aerobic condition. The increased growth rate in Δ*arcA* results from the higher uptake rate of oleate for its increased fitness. Thus, the faster the incorporation rate of oleate into the system, the faster the ^13^C dilution rate in amino acids.

Fluxes are governed by multiple factors, i.e. gene expression, posttranscriptional control and enzyme kinetics. *E.coli* WT grows sub-optimally in oleate metabolism as it exhibits broad substrate specificity by protein synthesis, suggesting the plasticity for the sudden shift from the glycolytic substrate to the gluconeogenesis substrate. Higher lag was encountered in WT under oleate but had little when grown in the preferred substrate (Basan et al., 2020).

Despite considerable knowledge of most aspects of fatty acid metabolism, the regulatory control of global transcription factors is still little understood. Our cost-effective strategy has displayed metabolic phenotypes of WT and Δ*arcA* by quantifying the dilution rates of position-specific isotopic distribution under fatty acid metabolism. Plausibly, this indicates that *E.coli* has evolved in a way where fatty acid has not remained a preferred carbon source. Enhanced ^13^C dilution rates along with the higher growth in Δ*arcA* potentially explains that the deletion has de-repressed the oleate uptake gene, β-oxidation pathway genes, along with the TCA cycle, and gluconeogenesis pathway. Thus, Δ*arcA* has manifested more flexibility to the system by its de-regulation. Therefore, this strain can act as a chassis in the industry for its enhanced ability to metabolize fatty acids by not compromising with the biomass yield. Furthermore, predictive modelling in getting fluxomics data will depict pathway fluxes, cofactor balance and metabolite accumulation.

## List of Abbreviation used

LCFA: long-chain fatty acids
arcA: aerobic respiration control protein
fad: fatty acid degradation
TCA: tricarboxylic acid
α-KG: alpha ketoglutarate
OAA: oxaloacetate
PEP: phosphoenolpyruvate
PYR: pyruvate
3PG: 3-phosphoglyceric acid
G6P: glucose-6-phosphate

## Author’s Contribution

S.J, K.V.V and S.K.M. conceived the idea, S.J and S.K.M designed the experiments, S.J. performed all the experiments. P.J assisted in the ^13^C GC-MS pipeline. S.J did the data mining. S.J, S.K.M analyzed the data. S.J, S.K.M, K.V.V wrote the original draft. S.J, S.K.M and K.V.V reviewed and edited the paper.

## Acknowledgments

We acknowledge the BioX Centre and Advanced Material Research Center of IIT Mandi for access to GC-MS, NMR and other facilities. S.J. thank the Council of Scientific and Industrial Research (CSIR), Government of India (09/087(0857)/2016-EMR-I) for the fellowship.

