## Supplemental 1 for "Oleate metabolism using kinetic ^13^C dilution strategy deciphered the potential role of global transcription regulator *arcA* in *Escherichia coli*"

**Sup 1: Strains Information**

Primer design information used in the present study

| **Name** | **Primer Sequence (5’-3’)** |
| --- | --- |
| **Primers used for generation of knockout** | |
| ArcA KO fwd | CTTTTGTACTTCCTGTTTCGATTTAGTTGGCAATTTAGGTAGCAAAC GTGTAGGCTGGAGCTGCTTCG |
| ArcA KO rev | CGGCGCTAAAAAGCGCCGTTTTTTTTGACGGTGGTAAAGCCGA CCGGGGATCCGTCGACC |
| **External primers (200 bp upstream and downstream to the gene) used for detection of knockouts** | |
| ArcA Locus Fwd | TTTTGACACTGTCGGGTCCTGAGGGAAAGT |
| ArcA Locus Rev | TTGGGAACCAGTGTGCTGGTGGTGG |

The E. coli variants used in the work are given as follows

| **Strain Name** | **Genotype** | **Source** |
| --- | --- | --- |
| *E. coli* K12 MG1655 WT | F-, λ-, rph- | Keio collection *CGSC #6300* |
| *E. coli* K12 MG1655 *Δ*arcA | F-, λ-, rph-, *ΔarcA kan* | This study |
